# Optimizing InterProScan representation generates a surprisingly good protein function prediction method

**DOI:** 10.1101/2022.08.10.503467

**Authors:** Henri Tiittanen, Liisa Holm, Petri Törönen

## Abstract

**Motivation:** Automated protein Function Prediction (AFP) is an intensively studied topic. Most of this research focuses on methods that combine multiple data sources, while fewer articles look for the most efficient ways to use a single data source. Therefore, we wanted to test how different preprocessing methods and classifiers would perform in the AFP task when we process the output from the InterProscan (IPS). Especially, we present novel preprocessing methods, less used classifiers and inclusion of species taxonomy. We also test classifier stacking for combining tested classifier results. Methods are tested with in-house data and CAFA3 competition evaluation data.

**Results:** We show that including IPS localisation and taxonomy to the data improves results. Also the stacking improves the performance. Surprisingly, our best performing methods outperformed all international CAFA3 competition participants in most tests. Altogether, the results show how preprocessing and classifier combinations are beneficial in the AFP task.

**Contact:** petri.toronen(AT)helsinki.fi

**Supplementary information:** Supplementary text is available at the project web site http://ekhidna2.biocenter.helsinki.fi/AFP/ and at the end of this document.

## 1 Introduction

As sequencing efficiencies are improving, post-processing and annotating the generated genomic and transcriptome sequences becomes more and more important [9, 16]. One of the major challenges is to annotate the large number of potential protein sequences. This has increased the importance of the Automated protein Function Prediction (AFP) in bioinformatics [15, 5]. AFP methods often collect data, usually from protein sequence, from interaction network and/or from gene expression data, and look for data features that can separate sequences with a certain function from the rest of the sequences. All the current AFP research, to our knowledge, compares the final prediction methods which often utilize multiple input data types. We see fewer articles, where just one input data is analyzed explicitly for optimal prediction accuracy. Such research can give strong recommendations for other method developers of how to use the discussed input more effectively.

InterProScan (IPS) features [13] are used frequently in the AFP methods, and many top-performers in recent CAFA competitions use IPS features as one of the inputs [26, 17, 11]. However, there is no publication that looks for the most efficient ways to use just IPS features in AFP task. Hence, we compare here different state-of-the-art classifiers and different preprocessing methods for IPS features. We also test the addition of extra features: Species taxonomy and the prediction of the protein cellular localization. Then, in order to get even better results, the predictions are combined with a second level classifier (classifier stacking [24, 20]), again also comparing different additional feature sets.

Our results have two levels of predictions that correspond to the two classification levels. Results from the first level show that it is beneficial to include taxonomy and the location of IPS features in the protein sequence to the data. Best performance was obtained here with rarely used classifiers. Next, the second level results show strong improvement from the first level. Finally, we compare our results against the latest AFP method competition, CAFA3 (Critical Assessment of protein Function Annotation [26]). Here our best second level classifiers outperform all the CAFA3 competition methods in Biological Process and Molecular Function with most metrics. This is quite a surprise, as our classifiers used just IPS features as input.

This article is organized as follows. We present the datasets, preprocessing steps, classifiers and the general classification scheme in Section 2.1 and 2.2. In Section 3, we describe the experiments and analyze the results. Finally, in Section 4, we conclude with a discussion of the results and present ideas for future work. Our supplementary text discusses the previous research and summarizes their IPS preprocessing methods and classifiers. It also discusses classifier methods and datasets in more detail.

## 2 Materials and methods

### 2.1 Datasets

We used two datasets in our experiment. Both contain IPS data as input features and GO data as predicted terms. We also refer to GO terms as GO classes, as this often simplifies the text. More detailed properties of the datasets are presented in the supplementary material.

#### 2.1.1 In-house GO data

Our in-house GO data is a collection of well-annotated proteins. This allows a thorough training and evaluation of the tested methods. The drawback is that the results cannot be compared to other published methods. For this, we use another dataset. Here, we collected GO annotations and sequence data from Uniprot from 2019 (date: 2019.10.16.).

Training and evaluating AFP methods requires reliable positive and negative data points for each class. However, this is difficult with biological annotations. We have only a subset of positive cases known and practically no confirmed negative cases. This is known as positive – unlabeled learning problem [2]. Therefore, the following steps aim to filter potential false positives and false negatives from our datasets.

In **the positive set**, we have a mixture of confirmed positive cases and unconfirmed predicted positive cases. These predictions were created by UniProt. Here we selected only clear manually curated annotations, and excluded GO evidence codes IEA (Inferred from Electronic Annotation) and ND (Not Defined). We also excluded annotations based on sequence similarity (ISS, Inferred from Sequence Similarity) as these might generate circular logic while selecting sequence features for annotations. This selects the well confirmed positive datapoints.

Furthermore, some positive predictions are linked to very vague functional classes in the GO hierarchy. These are harmful to training and evaluation, as they add false negative cases to deeper precise classes in our datasets. Here, we selected only classes that had class size smaller than 5% from the root node of the ontology. In addition, we excluded direct annotations to some selected GO classes that, although not too small by our size criteria, were still known to be too vague for presenting any true information. These GO classes were classes like ‘Protein Binding’. We have used similar procedure earlier in Törönen et al. [22]. Notice that we still allow annotations to large classes in our datasets, when they are parental (ancestral) classes to more specific classes in GO structure. This selects the annotations that are specific enough to our dataset.

For **the negative set**, we select the genes that have some other positive annotation, as described above, while lacking the annotations to predicted or evaluated GO class. We estimate that this filters potential class members from the unlabeled pool. Defining positive and negative data points for AFP task is an open research question, and there are alternative solutions [7, 25].

#### 2.1.2 CAFA3 GO data

CAFA3 analysis uses separate training and evaluation datasets. The evaluation dataset is distributed by CAFA organizers [26]. The CAFA3 evaluation data is smaller and its annotations are less detailed, when compared to our in-house data. However, it allows the comparison against the CAFA3 competition participants. The training dataset was generated in similar fashion as our in-house training dataset. The difference was that we used the GO data collected from 11.29.2016. We also used the sequence data and IPS version from the same time. Otherwise, exactly the same filtering procedures were applied here as with in-house data.

For the evaluation part, we use the data collected during the CAFA3 accumulation period [26]. This contains sequences, not annotated in the database during the training period. This data gives us a reference point against the state-of the art methods. We used only sequences with no previous knowledge (No Knowledge data) here.

#### 2.1.3 Sequence-based features

InterProScan (IPS) output [13] was the main component of input data. IPS scans query protein for various protein domains, sequence motifs and protein families, by using several related databases. For in-house data, we used the most up-to-date version of IPS at the time (5.38-76.0). For the CAFA dataset, we selected the IPS version that was available at the CAFA3 training time (5.22-61.0).

We also added predictions for cellular location to our input data. These are usually not as detailed predictions as what Cellular Component in GO data stores. Still, we considered these as potentially useful information for the prediction tasks. Here we used TargetP [4] and WolfPSort [12] programs. Figure 1 shows an overview how different features are created in our project.

**Figure 1:**
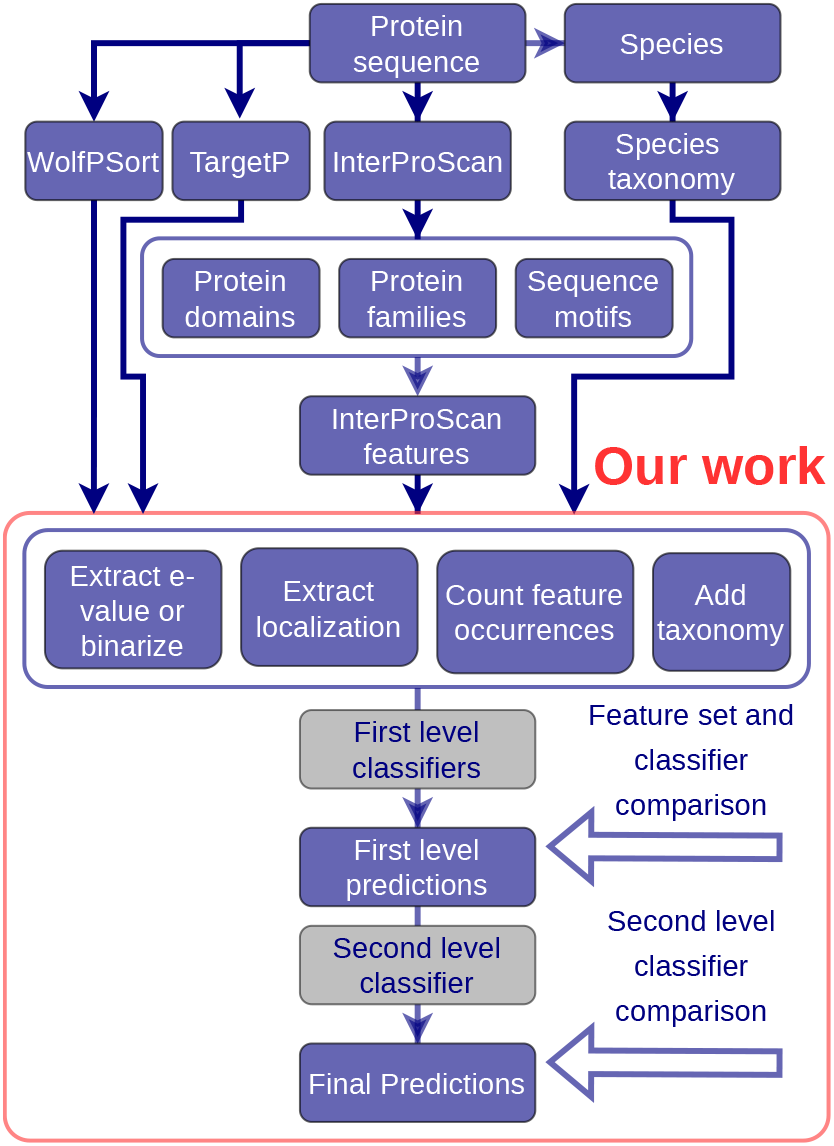
Schematic workflow of the classification process. A query protein is sent to InterProScan, which extracts various sequence features from it. We also extract species information from sequence and map it to NCBI species taxonomy. We also predicted the cell compartment for sequences with SignalP and WolfPSort. Next, we processed the InterProScan features in various ways and tested each first level classifier with each feature set separately. Final step tests different second level classifiers for combining first level classifiers.

#### 2.1.4 Species taxonomy features

We also tested the addition of species taxonomy to our input feature set. Here the aim is to allow a classifier to generate different predictions with same sequence feature when the sequence occurs in different regions of species taxonomy tree. We were the first research group to use taxonomy in our AFP method [14], and it has been since used, to our knowledge, by only one other research group [11]. Here we used a script (from [22]) that takes the species taxonomy identifier, maps it to NCBI taxonomy hierarchy and links species to its taxonomic groups. This taxonomic group list is finally converted to binary vector.

#### 2.1.5 IPS Feature sets

One of our main goals was to find efficient ways to process IPS output. For this, we constructed different IPS based feature sets for both in-house and CAFA3 data. We were especially looking at the following questions:

1. Is the classifier performance better with binarized or with continuous IPS feature data?
2. Is it beneficial to include external information, like species taxonomy, to the prediction process?
3. Should we extract additional information from the IPS features?

For **question 1** we are converting IPS features either to binary vectors or converting the associated E-values with negative log-function (see Table 1). These are our *binary* and *e-value* feature set. For **question 2** we combine the selected IPS feature set, either *e-value* or *binary*, with the binary taxonomic vector. This is our *taxonomy* feature set.

**Table 1:** Explanation of different feature sets. How the feature set were generated. What specific question is studied with the feature set.

| Feature Set | Method | Research question |
| --- | --- | --- |
| *e-value* | Negative log transformed IPS features | Does the inclusion of IPS feature strength improve prediction? |
| *binary* | All non-negative log-evalues are converted to one | Is binary score better than the weighted version? |
| *cluster* | Include IPS feature clusters to feature set | Does the inclusion of feature clusters improve the prediction? |
| *taxonomy* | Add taxonomy features to selected feature set | Does the addition of taxonomy features improve the prediction result? |
| *count* | Include also the count value of each feature to feature set | Is the number of feature occurrences useful for the prediction? |
| *location* | Include the location of the feature to feature set | Is the location of the feature useful for the prediction? |

For **question 3** we tested three alternatives. First, we include the *rough sequence location of the feature* to our feature set. Here the sequence is divided evenly into three equally sized parts: start, middle and end. Three location features are constructed for each IPS feature based on that split, each containing the proportion of the feature in the corresponding part of the sequence. This is our *location* feature set. Our second alternative is to use *IPS feature clusters* as additional features. These clusters group together features that are closely related to each other. Here we select the strongest signal, from the features within the IPS feature cluster, to be the signal for the specific cluster. Finally, we also test the *count information* for each IPS feature. Here, we simply count how many times each IPS feature occurs in the sequence. All the discussed feature sets are summarized in supplementary text

Finally, the feature sets *cluster*, *location*, *count* and *taxonomy* are always constructed by combining the respective features with the base features i.e. *e-value* or *binary*. The resulting quite high dimensional feature sets, shown in supplementary text, allow classifiers to learn more complex models and extract signals that cannot be detected from the simpler feature sets. Unfortunately, the larger feature dimensions can potentially dilute the signal that is strong in a more compact feature set and make the search of relevant features totally unfeasible for many algorithms.

#### 2.1.6 Additional features in stacking process

We also investigate the effect of combining predictions for increased accuracy, i.e. second level prediction (classifier stacking). Similarly to the first level experiments, we evaluate a set of additional feature sets that are here used in combination with first level predictions as inputs to the second level classifiers. We evaluate combinations of first level predictions, first level predictions converted to rank values, *taxonomy* and a set of *additional* features consisting of the following features: sequence length, the number of X letters in the whole sequence, the number of IPS features found, the proportion of the area covered by the IPS features in the whole sequence and the signal of the strongest IPS feature. Selected additional features measure the overall quality of signal of IPS features.

### 2.2 Classification Outline

The general classification scheme is presented in Figure 1. The classification is done in two stages. First, a set of different classifiers are trained and evaluated on different feature set combinations of IPS data. This shows the effect of different feature sets and different classifiers to the classification performance. Next, second level classifiers are trained and evaluated on the selected first level predictions of different classifier-feature set combinations. Here we also test additional features at the second level. Finally, the second level results are compared to the first level predictions to evaluate the effectiveness of model stacking and additional features. A separate classifier is trained for each GO class in both first and second stage.

We use model stacking [24], a widely studied method [20], for combining the first level predictions. Model stacking is an ensemble learning method where predictions of multiple classifiers are used as input to a second level classifier, with the aim of achieving better performance than any single first level classifier.

Our experiments with the in-house data are evaluated using cross validation. Although the details of cross validation are later in Cross Validation section 2.3.1, we cover here how it is used in stacking. Specifically, all first level classifiers are first trained with training data 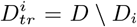 and then used to predict the results for evaluation data 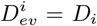. This process is repeated, until predictions are created for every *D_i_*. Next, the second level classifiers are trained with the first level classifier predictions, over the data points in 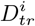, to generate the predictions for 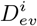. This is again repeated for every value of *i*. A similar principle can be used to train third level classifiers.

The combination of cross validation and separate classifiers for each GO class requires that we train five classifiers for each GO class. This leads to massive number of models with each classifier (CC: 8440, MF: 17240 and BP: 56440, with in-house data). Therefore, in order to obtain results in a reasonable time, it was necessary to limit the training time of individual classifiers to around eight minutes for a single class. Thus, it would be possible to get better results for individual classes than presented here if more training resources were used. One of the benefits of the class specific training is that it would make it possible to easily select the time used to train a classifier for a particular class.

#### 2.2.1 First Level Classifiers

We selected a set of classifiers with different operating principles e.g. linear, nonlinear and tree like to be compared in the first level classification experiments. They produce different types of decision surfaces, and therefore provide a good basis for stacking [24]. An overview of the used classifiers is presented by Table 2 in the supplementary text. We included classifiers that have been used widely in the AFP domain, such as basic logistic regression (LR), support vector machine (SVM), and neural network (ANN). We also tested classifiers that have been used very little in the AFP research, like extreme gradient boosting (XGB) and factorization machine (FM). To our knowledge, FM has not been used previously in the AFP domain, and only one use of XGB was found just when we were finishing this article [23].

**Table 2:** First level cross validated classifier performance (AUC-PR, larger value is better) on in-house data with different feature sets. The highest score for each classifier is bolded.

| Data | Classifier | *e-value* | *binary* | *e-value*<br>+ *cluster* | *binary*<br>+ *cluster* | *location* | *count* | *taxonomy* |
| --- | --- | --- | --- | --- | --- | --- | --- | --- |
| CC | XGB | 0.677 | 0.654 | 0.678 | 0.656 | 0.702 | 0.690 | **0.723** |
|  | E.NET | 0.518 | 0.394 | **0.520** | 0.396 | 0.518 | 0.519 | 0.513 |
|  | SVM | 0.477 | 0.520 | 0.478 | 0.524 | **0.552** | 0.528 | 0.544 |
|  | FM | 0.143 | 0.574 | 0.143 | 0.578 | **0.654** | 0.628 | 0.629 |
|  | ANN | 0.538 | 0.546 | 0.525 | **0.556** | 0.222 | 0.555 | 0.546 |
| MF | XGB | 0.822 | 0.778 | 0.823 | 0.782 | 0.844 | 0.832 | **0.874** |
|  | E.NET | 0.654 | 0.544 | **0.655** | 0.547 | 0.654 | 0.654 | 0.649 |
|  | SVM | 0.651 | 0.681 | 0.652 | 0.686 | **0.705** | 0.687 | 0.704 |
|  | FM | 0.135 | 0.738 | 0.143 | 0.742 | **0.785** | 0.768 | 0.744 |
| BP | XGB | 0.550 | 0.515 | 0.550 | 0.517 | 0.572 | 0.561 | **0.597** |
|  | E.NET | **0.365** | 0.271 | 0.356 | 0.273 | 0.355 | 0.351 | 0.352 |
|  | SVM | 0.336 | 0.373 | 0.336 | 0.376 | 0.409 | 0.391 | **0.415** |
|  | FM | 0.085 | 0.432 | 0.143 | 0.436 | 0.507 | 0.479 | **0.525** |

As a linear model, we used logistic regression with elasticnet loss (e.net) [27], which is suitable for our high dimensional data due to its sparsity and robustness. Support vector machine was used with a radial basis function kernel. Support vector machines are a popular choice for high dimensional data. However, their memory consumption is prohibitively high for datasets of the size used here. Hence, we had to down sample the data to make them operational. The neural network used was small with two hidden layers of size 5, rectified linear unit (ReLu) activation function, batch size 100 and 3 iterations. The computation time limits imposed by the high number of classes, and the presence of very small classes are not an ideal situation for artificial neural networks. The small ANN size and low amount of iterations was necessary because of the training time constraints. Note that we currently omitted ANN from first level, as it required considerable running times in our preliminary test. Extreme gradient boosting [6], is a powerful and widely used classifier, based on classification trees. The factorization machine [19] is a relatively new and less used classifier, that is related to matrix factorization models and support vector machines. It is very scalable and fast for large scale sparse data and therefore very suitable for AFP task. Note again that we are limited here to fast and well scaling classifiers.

Our current work excludes popular Deep Neural Networks (DNN). We were forced to this for the following reasons: a) Our current Cross Validation step was done separately for each predicted GO class. This would have required an enormous number of separate DNNs. b) Also, the benefits of DNNs are lost when we train one class instead of training multiple classes. c) Many classes had quite few positive samples in our training data. This would have been challenging for training DNN methods.

#### 2.2.2 Second Level Classifiers

The first level classifier selection was constrained by the very high dimensional data. The dimensionality of the second level data is low compared to the first level data. Hence, the classifiers used in the stacking phase are different or readjusted. The classifiers used in the second level are XGB, LR, XGB-ltr (XGB with learning to rank loss) and ANN. The training time of individual classifiers is still limited, which is taken into account in model selection and parameterization, as explained below. The relatively low dimensional data made it possible to use non-sparse losses for logistic regression. Hence, we selected logistic regression with the standard l2 loss. Gradient boosting was used with modified parameters. Most notably, the number of trees was increased. We also tested XGB also with a pairwise learning to rank loss function. The second level ANN architecture was with one hidden layer of size 5, ReLU activation and batch size 100.

We used predictions from XGB, SVM, FM and e.net, where each classifier was trained on *location* and *taxonomy* feature sets separately, as inputs. Therefore, the second level classifiers had total eight separate first level predictions among the input features. These feature sets were chosen because their good performance in the first level experiments. Here, for SVM, ANN and LR the features were preprocessed by removing the mean and scaling to unit variance.

When testing the methods on CAFA3 data, we also used the following baseline stacking approaches to compare the performance of the more complex methods presented above: Mean, which is the mean of the first level predictions and r.mean which is the rank averaged mean of the first level predictions.

### 2.3 Method Evaluations

#### 2.3.1 Cross validation with in-house data

The experiments with the in-house data were evaluated using cross validation. We used here stratified cross validation that was performed separately for each GO class. This enables reliable training and evaluation of very small classes. Our Supplementary text discusses this topic more in detail. Unfortunately it excludes comparisons with many existing AFP methods and also Protein-Centric evaluation (explained in Evaluation Metrics 2.3.3). However, it should be noted that our tests with CAFA3 evaluation datasets represent a very detailed protein-centric evaluation with four evaluation metrics. Finally, we point that separate stratified cross validation for each GO class is problematic, and our Discussion 4 points our future directions to solve this.

#### 2.3.2 Evaluation with CAFA data

Experiments with CAFA3 data started similarly to our analysis with inhouse data. First, we create the class-specific cross validation splits. Next, for each fold in a split, we train the first level classifiers on the remaining folds and get predictions for the current fold. We combine the first level predictions and train a second level classifier on the predictions, as described in section Classification Outline. Next, the processing differs from the in-house dataset. We first retrain the first-level classifiers using the whole data. Next, we generate predictions for CAFA evaluation dataset with first level classifiers. Finally, we used the previously learned second level classifiers to combine the first level predictions for CAFA evaluation set. Here we have a completely separate evaluation dataset from the training process. Also, this data allows both protein-centric and term-centric evaluations. CAFA evaluation used No Knowledge data (NK-data) with ALL setting.

#### 2.3.3 Evaluation metrics

Selecting evaluation metrics for AFP is a difficult and often overlooked task [18, 15]. Still, it has a drastic impact on results, and some popular evaluation metrics are not well suited for AFP task [18, 3, 8]. Here we used metrics that are either well suited to AFP evaluation or allow comparison against the latest CAFA competition. Here we summarize the used evaluation metrics and give a more detailed description in our supplementary text.

The used evaluation metrics form two groups: *Term Centric* (TC) and *Protein Centric* (PrC) evaluation metrics. TC metrics process each GO class (GO term) separately and summarize the results. They are insensitive against biases generated by different class sizes, but require often more data to work well (see supplementary text). PrC metrics process each protein separately and combine these results. They are often sensitive to biases between class sizes [18] but can work with smaller amount of data. For our cross validation evaluation we use Term Centric Area Under the Precision Recall Curve (TC-AUCPR). For our CAFA evaluation we used all five CAFA evaluation metrics: maximum of F-score (*F_max_*), minimum of semantic similarity (*S_min_*), normalized semantic similarity *nS_min_*, weighted F-score (wF_max_) and TC Area under area under Receiver Operating Characteristics curve (TC-AUC). All these measures, except for TC-AUC, are PrC metrics.

Using these five evaluation metrics in parallel has its benefits: Each metric is expected to have its own biases and errors. These weaknesses are lessened when we monitor five metrics in parallel. Furthermore, we and others have shown that *F_max_*, is a biased metric [18, 8, 3]. Here the inclusion of other evaluation metrics, *S_min_, nS_min_*, wF_max_ and TC-AUC allows us to check if they can generate a more reliable consensus.

### 2.4 Visual Analysis of First level classifiers

As we generated a separate score, AUC-PR, for each classifier with each GO class, it is interesting to check how different classifier - GO class combinations perform. We visualized this with heatmap function from R environment. Heatmap creates a matrix visualization, where number values are presented with color. Block structures are revealed by ordering rows and columns with hierarchical clustering. Cluster trees were further rearranged, for better visualization, with a function from R Seriation package [10]. Rearranging is explained more here [1].

## 3 Results

### 3.1 First-level Classifiers with In-house data

The first level cross validated classification results for the in-house data are presented in table 2. We first compared *e-value* and *binary* with each other. Here it is clear that the optimal choice between the base features *e-value* and *binary* depends mainly on the properties of the classifier. FM, for example, is designed for binary data, showing good performance only on binary data. SVM shows similar but weaker difference, whereas XGB and e.net show the opposite behavior (see table 2).

Next we test additional IPS information. These are mainly tested with the better of two first feature sets. *cluster* gave here only minimal increase in the scores at best. Next, *location* is beneficial for nearly all classifiers, often being the best feature set. *count* usually improves classification performance, but not as much as *location*. *taxonomy* gives very high increase for XGB but does not give as significant improvement to the other classifiers. XGB, as a tree based model, might be better at combining non-linear signals here.

Comparing classifiers shows that XGB is the best performing classifier for all ontologies and especially benefits from taxonomy information. FM is generally the second best. SVM and e.net results are far from XGB and FM results. The big size of the datasets and the extremely large dimensionality potentially made it hard to get SVM to perform well. The nonlinear classifiers FM and XGB seemed to give better results than linear e.net. Although the ANN used was very small, its training time resulted to be prohibitively slow for other datasets than CC and thus ANN results for other datasets are omitted.

### 3.2 Visual analysis of First-level Classifiers

We created heatmap visualizations to classifiers at single class level. This looks if the performance order of classifiers is the same across all the classes. Results are shown in figure 2. We can clearly see, that the classifiers often form distinct blocks where a particular group of classifier performs well. This is a clear indication that a class specific combination of the classifier predictions has chances to outperform any single classifier.

**Figure 2:**
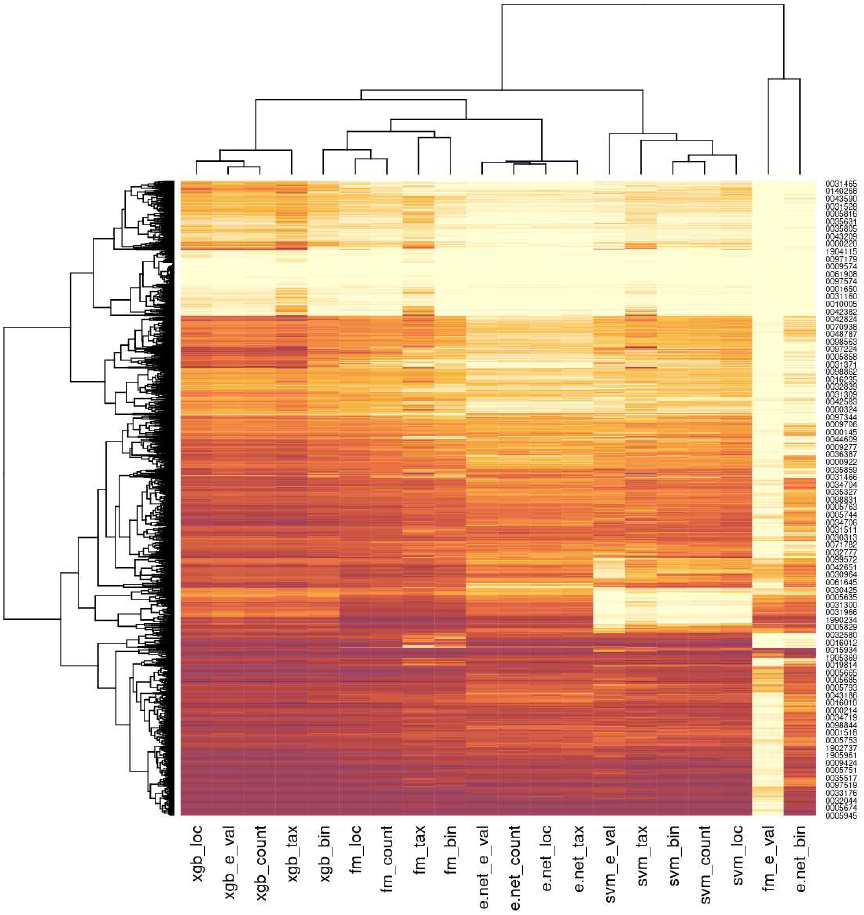
Class specific AUC-PR classification performance (darker color is better) of a set of classifiers (columns) for a set of classes (rows). This figure shows results for predicted Cellular Component GO classes from in-house data. Different classes have different best performing classifiers.

The heatmap has some interesting groups of GO classes, but we discuss them here only shortly. First, the classifiers form roughly four clusters to columns: XGB, FM, e.net and SVM. Two weak performing clusters form an outlier cluster (FM with *e-value* and e.net with *binary*). The very bottom of the heatmap shows GO classes that are predicted well by most methods. This is also a region where we see no clear difference between prediction methods. The most striking cluster of the heatmap is the cluster approximately one third away from the bottom. It contains GO class cluster where SVM fails completely and FM classifiers are the best performers. Very top shows a GO class cluster where XGB classifiers are slightly better from other methods. Below it, we see GO classes where all classifiers fail. Only three classifiers, XGB, FM and SVM with *taxonomy* feature set show some performance at lower regions of this cluster. Altogether, this figure suggests that: A) Best classifiers and feature sets can vary significantly across the predicted classes, like here with GO classes. B) Evaluating classifiers without looking at the class-level performance can oversimplify the differences between classifiers.

### 3.3 Second-level Classifiers with In-house data

The second level classification results for the in-house data are presented in Table 5. Feature set comparison shows that *taxonomy* features give the best results for all top performing classifiers and the biggest increase from the base level. Although *taxonomy* features were present already in the first level, the information contained in them is still very useful in the second level. *additional* features marginally increase performance, and since it is a very low dimensional feature set, including it is recommended. Rank features show minimal benefit for ANN, but they do not seem to give any benefit for other classifiers.

Comparing results by classifier shows that XGB gives again the best scores followed by LR while XGB-ltr and ANN performance is generally lower. The base models are far behind the main classifiers. From the base models, mean stacking gives small improvement for CC and BP over the first level, still losing to the best stacking methods. For MF, mean stacking does not outperform the best first level result. It is likely that some of the low quality MF first level classifier predictions may affect the mean too much here.

Comparing Tables 2 and 5, we can see that stacking improves the best result for CC and BP, but for MF there is no improvement. The first level scores for MF are potentially already so high that they cannot be improvement. In general, stacking can be concluded to be beneficial.

### 3.4 First-level Classifiers with CAFA data

The CAFA3 data first level results are presented in Table 3. We limit this analysis only to basic IPS (*binary* or *e-value*), *taxonomy* and *location* feature sets. Otherwise, the result table would be massive. Here, *taxonomy* and *location* feature sets give often again an increase in performance compared to basic IPS feature data. Again, XGB and FM are the top performing classifiers, but now FM is the best one, often with a clear margin to XGB. In MF, e.net results are surprisingly good. In general, these CAFA3 comparison results seem to be in accordance with the in-house data results presented earlier.

**Table 3:**
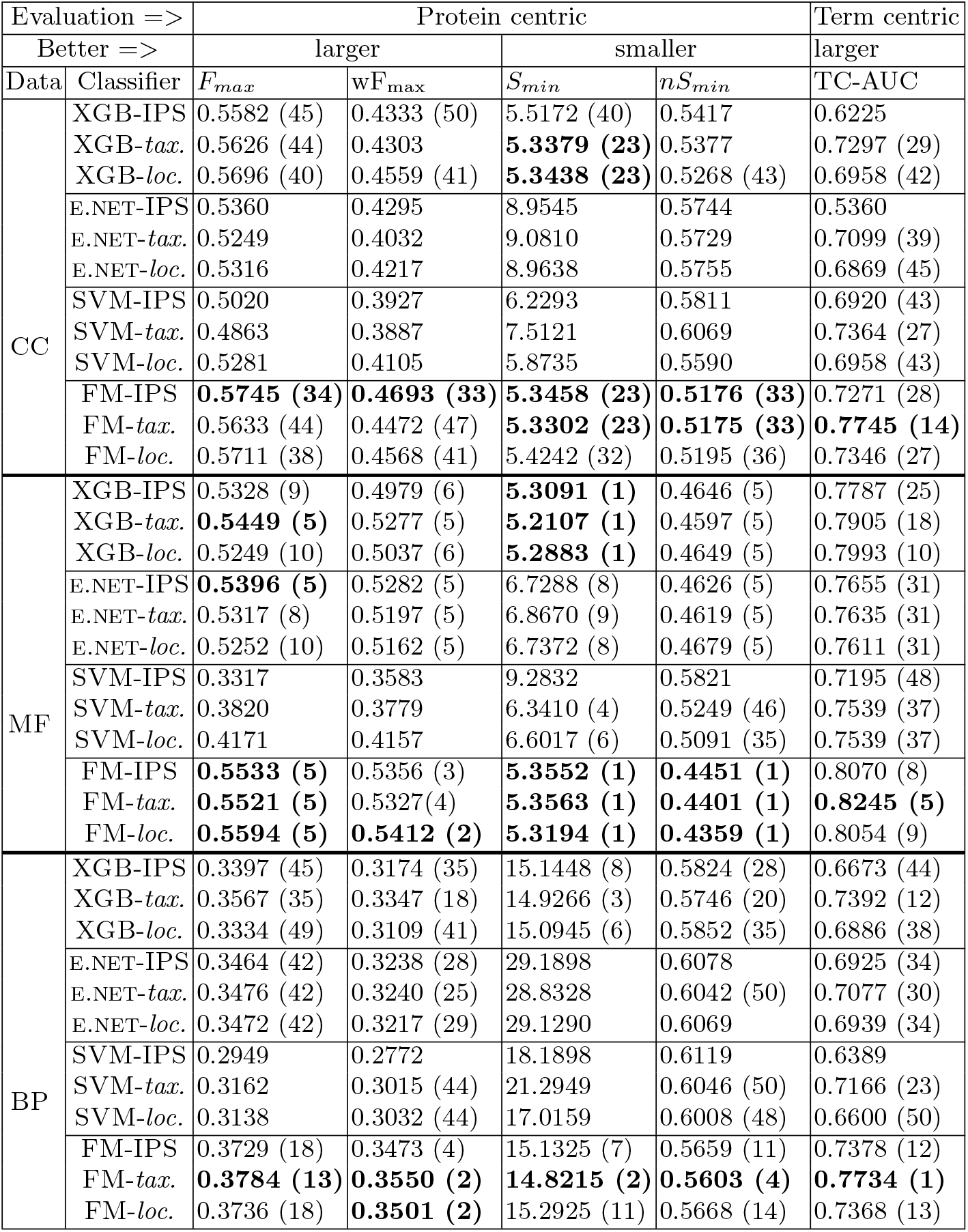
CAFA3 first level results. We compare results using all reported CAFA3 evaluation metrics. We also include *F_max_* into our comparison, although it has been shown to be very biased evaluation metric [18, 3, 8]. FM and XGB show best performance, matching the CAFA3 top methods in BP and MF. We exclude the ranking, if the method is outside the top-50.

In addition to the evaluation metric scores, we also show rank of each score against CAFA3 competition results. Rank value is omitted if the score is not among top-50 results. The best MF an BP rankings are here really high. Especially XGB and FM outperform all CAFA3 methods in *S_min_* metric. Other metrics, however, suggest lower ranking, although still mostly within top-10. Still, the rankings vary quite a lot between the metrics. This underlines the relevance of using a group of evaluation metrics in parallel.

### 3.5 Second-level Classifiers with CAFA data

Results from the second level classification experiments performed on the CAFA3 data are presented in Table 4, and a summary of the first and second level results is presented in Table 6. We can see, that the stacking results are again better than the first level results. However, the improvement is smaller compared to the stacking improvements observed with the in-house data. Here, the different evaluation metrics *F_max_*, wF_max_, *nS_min_*, TC-AUC and *S_min_* rank the methods in different order. XGB and ANN are generally the strongest classifiers. LR performance is relatively very low for MF compared to CC and BP, while XGB stacking is still better than the first level XGB on MF. Therefore, MF may be more nonlinear in nature than CC and BP. The small ANN performs better on stacking level than as a first level method, since the lower dimensionality of the stacking problem is easier to optimize in the limited training time.

**Table 4:** CAFA3 second level results. Brackets show again the ranking against the CAFA3 competition methods. Again, we include also *F_max_* to comparison although it is a biased metric (see table 7). Notice that we generate top-ranking results with XGB and ANN in BP and MF. Ranking in CC varies heavily between metrics.

| Evaluation => |  | Protein centric |  |  |  | Term centric |
| --- | --- | --- | --- | --- | --- | --- |
| Better => |  | larger |  | smaller |  | larger |
| Data | Classifier | $F_{max}$ | wF<sub>max</sub> | $S_{min}$ | $nS_{min}$ | TC-AUC |
| CC | LR | 0.5647 (43) | 0.4029 | 5.3473 (23) | 0.5501 | 0.5592 |
|  | XGB | 0.5782 (30) | 0.4666 (37) | 5.1437 (3) | 0.5062 (24) | **0.8161 (4)** |
|  | ANN | 0.5670 (40) | 0.4402 (50) | 5.4205 (28) | 0.5300 (47) | 0.7262 (28) |
|  | MEAN | 0.5796 (30) | 0.4514 (44) | 5.1668 (5) | 0.5126 (32) | 0.8029 (5) |
|  | R.MEAN | **0.5988 (10)** | **0.4810 (20)** | **4.9481 (1)** | **0.4975 (19)** | 0.8082 (5) |
|  | LTR-XGB | 0.5140 | 0.4085 | 5.5147 (40) | 0.5457 | 0.8079 (5) |
| MF | LR | 0.3419 | 0.3374 | 10.2349 | 0.6236 | 0.7378 (41) |
|  | XGB | **0.5819 (5)** | **0.5660 (1)** | **4.9658 (1)** | **0.4198 (1)** | 0.8331 (4) |
|  | ANN | **0.5641 (5)** | **0.5470 (1)** | **5.1194 (1)** | **0.4325 (1)** | 0.8255 (5) |
|  | MEAN | **0.5596 (5)** | 0.5410 (2) | **4.9603 (1)** | **0.4266 (1)** | **0.8412 (3)** |
|  | R.MEAN | 0.4373 | 0.4335 | 7.2234 (21) | 0.5094 (35) | 0.7715 (29) |
|  | LTR-XGB | 0.4693 | 0.4684 (30) | **5.4749 (1)** | 0.4882 (9) | 0.8207 (7) |
| BP | LR | 0.3716 (19) | 0.3392 (6) | 14.8407 (3) | 0.5675 (13) | 0.6810 (41) |
|  | XGB | **0.3888 (3)** | **0.3656 (1)** | **14.6046 (1)** | **0.5473 (1)** | **0.7887 (1)** |
|  | ANN | 0.3846 (7) | **0.3578 (1)** | **14.4483 (1)** | 0.5534 (2) | **0.7872 (1)** |
|  | MEAN | 0.3799 (12) | **0.3530 (1)** | **14.5686 (1)** | 0.5531 (2) | **0.7737 (1)** |
|  | R.MEAN | 0.3583 (34) | 0.3365 (15) | 15.1847 (8) | 0.5716 (20) | 0.7697 (3) |
|  | LTR-XGB | 0.3210 | 0.3076 (44) | 15.1159 (8) | 0.5870 (37) | 0.7726 (2) |

**Table 5:** Second level cross validated classifier performance (AUC-PR, larger value is better) on in-house data with different feature sets. The highest score for each classifier is bolded. Simple baseline classifier performance is included for comparison.

| Data | Classifier | First level | First level<br>+ rankings | First level<br>+ *additional* | First level<br>+ *additional*<br>+ rankings | First level<br>+ *additional*<br>+ *taxonomy* |
| --- | --- | --- | --- | --- | --- | --- |
| CC | XGB | 0.769 | 0.766 | 0.776 | 0.772 | **0.781** |
|  | LR | 0.758 | 0.757 | 0.757 | 0.756 | **0.760** |
|  | ANN | 0.753 | **0.756** | **0.756** | 0.755 | 0.744 |
|  | XGB-ltr | 0.752 | 0.738 | 0.759 | 0.747 | **0.764** |
| MF | XGB | 0.866 | 0.865 | 0.870 | 0.868 | **0.873** |
|  | LR | 0.867 | 0.866 | 0.866 | 0.867 | **0.869** |
|  | ANN | 0.857 | **0.860** | 0.859 | 0.859 | 0.857 |
|  | XGB-ltr | 0.848 | 0.840 | 0.854 | 0.846 | **0.857** |
| BP | XGB | 0.642 | 0.638 | 0.649 | 0.644 | **0.655** |
|  | LR | 0.633 | 0.631 | 0.630 | 0.628 | **0.638** |
|  | ANN | 0.623 | **0.629** | 0.628 | 0.628 | 0.612 |
|  | XGB-ltr | 0.619 | 0.596 | 0.627 | 0.606 | **0.630** |
| Simple baseline stacking methods |  |  |  |  |  |  |
| Data | Classifier |  | Score |  |  |  |
| CC | Mean |  | 0.741 |  |  |  |
|  | Ranking Mean |  | 0.642 |  |  |  |
| MF | Mean |  | 0.854 |  |  |  |
|  | Ranking Mean |  | 0.741 |  |  |  |
| BP | Mean |  | 0.602 |  |  |  |
|  | Ranking Mean |  | 0.502 |  |  |  |

**Table 6:** Final comparison against the CAFA3. We show best first and second level methods against previously used classifier - feature set combinations. Best classifiers include any classifier that obtains a top score in any of the monitored evaluation metrics. Previously used classifiers represent e.net and SVM without the proposed added features. Again, the brackets show the ranking in the CAFA3 competition and *F_max_* is included, although it is a biased metric. The column Lvl shows the classification level. Notice how rankings improve drastically as we move from basic methods to best methods at the second level. XGB from second level shows overall best performance in MF and BP and shows overall best performance in CC.

| Data | Lvl | Method | $F_{max}$ | wF<sub>max</sub> | $S_{min}$ | nSmin | TC-AUC |
| --- | --- | --- | --- | --- | --- | --- | --- |
| CC | 1 | E.NET-IPS | 0.5360 | 0.4295 | 8.9545 | 0.5744 | 0.5360 |
|  | 1 | SVM-IPS | 0.5020 | 0.3927 | 6.2293 | 0.5811 | 0.6920 (43) |
|  | 1 | XGB-*tax.* | 0.5626 (44) | 0.4303 | 5.3379 (23) | 0.5377 | 0.7297 (29) |
|  | 1 | XGB-*loc.* | 0.5696 (40) | 0.4559 (41) | 5.3438 (23) | 0.5268 (43) | 0.6958 (42) |
|  | 1 | FM-IPS | 0.5745 (34) | 0.4693 (33) | 5.3458 (23) | 0.5176 (33) | 0.7271 (28) |
|  | 1 | FM-*tax.* | 0.5633 (44) | 0.4472 (47) | 5.3302 (23) | 0.5175 (33) | 0.7745 (14) |
|  | 2 | XGB | 0.5782 (30) | 0.4666 (37) | 5.1437 (3) | 0.5062 (24) | **0.8161 (4)** |
|  | 2 | R.MEAN | **0.5988 (10)** | **0.4810 (20)** | **4.9481 (1)** | **0.4975 (19)** | 0.8082 (5) |
| MF | 1 | E.NET-IPS | **0.5396 (5)** | 0.5282 (5) | 6.7288 (8) | 0.4626 (5) | 0.7655 (31) |
|  | 1 | SVM-IPS | 0.3317 | 0.3583 | 9.2832 | 0.5821 | 0.7195 (48) |
|  | 1 | XGB-*tax.* | **0.5449 (5)** | 0.5277 (5) | **5.2107 (1)** | 0.4597 (5) | 0.7905 (18) |
|  | 1 | FM-*tax.* | **0.5521 (5)** | 0.5327 (4) | **5.3563 (1)** | **0.4401 (1)** | 0.8245 (5) |
|  | 1 | FM-*loc.* | **0.5594 (5)** | 0.5412 (2) | **5.3194 (1)** | **0.4359 (1)** | 0.8054 (9) |
|  | 2 | XGB | **0.5819 (5)** | **0.5660 (1)** | **4.9658 (1)** | **0.4198 (1)** | 0.8331 (4) |
|  | 2 | ANN | **0.5641 (5)** | **0.5470 (1)** | **5.1194 (1)** | **0.4325 (1)** | 0.8255 (5) |
|  | 2 | MEAN | **0.5596 (5)** | 0.5410 (2) | **4.9603 (1)** | **0.4266 (1)** | **0.8412 (3)** |
| BP | 1 | E.NET-IPS | 0.3464 (42) | 0.3238 (28) | 29.1898 | 0.6078 | 0.6925 (34) |
|  | 1 | SVM-IPS | 0.2949 | 0.2772 | 18.1898 | 0.6119 | 0.6389 |
|  | 1 | FM-*tax.* | 0.3784 (13) | 0.3550 (2) | 14.8215 (2) | 0.5603 (4) | **0.7734 (1)** |
|  | 1 | FM-*loc.* | 0.3736 (18) | 0.3501 (2) | 15.2925 (11) | 0.5668 (14) | 0.7368 (13) |
|  | 2 | XGB | **0.3888 (3)** | **0.3656 (1)** | **14.6046 (1)** | **0.5473 (1)** | **0.7887 (1)** |
|  | 2 | ANN | 0.3846 (7) | **0.3578 (1)** | **14.4483 (1)** | 0.5534 (2) | **0.7872 (1)** |
|  | 2 | MEAN | 0.3799 (12) | **0.3530 (1)** | **14.5686 (1)** | 0.5531 (2) | **0.7737 (1)** |

The biggest surprise in second level results is that our top performing stacking methods, XGB and ANN, outperform all the CAFA3 methods in MF, while using evaluation metric *S_min_*, *nS_min_* or wF_max_. Also the ranks from the two remaining metrics, TC-AUC and *F_max_*, suggest that we would have been top-5. Results for BP are even better, as there ANN is the best in wF_max_, *S_min_* and TC-AUC and the second best in *nS_min_*. XGB further tops this by outperforming all CAFA3 methods in wFmax, *S_min_, nS_min_* and TC-AUC.

Another equally big surprise is how inconclusive and scattered the CC results are, when different evaluation metrics are viewed. XGB, the top-performer in BP and MF, is ranked as third in *S_min_* and fourth in TC-AUC. The other metrics, however, rank XGB here between 37*^th^* and 24*^th^*. Mean is ranked fifth by *S_min_* and TC-AUC, but also ranked low by other evaluation metrics. r.mean CC results are surprisingly good. It is ranked as best by *S_min_* and fifth by TC-AUC. Still also its rankings, when other metrics are used, are clearly lower. Notice that r.mean is quite rough classifier stacking method. Altogether r.mean results seems to be caused by the characteristics of CC, since its MF and BP results are not generally among the best.

As a summary, these results suggest that different second level methods would generate the best CAFA3 results in different subsets of GO. Biggest difference is seen between CC and other two subsets. We argue in discussion that this is probably caused by some weaknesses in CC evaluation data.

## 4 Discussion

### 4.1 Classifiers and feature sets

We have presented a comparison of different ways to use IPS features with various types of classification models on two datasets and diverse evaluation measures. This is the only article, to our knowledge, that has tested different ways to process IPS output. Our results from in-house dataset show that:

- Some classifiers benefit from inclusion of E-value to input data (XGB and e.net)
- Taxonomy and IPS location based features improve results over basic IPS features.
- Less used classification methods XGB and FM give the best performance at the first level.
- Our visual analysis shows that top performing classifiers vary clearly across the GO classes.
- Class specific second level stacking improves the results, often with a clear margin.

Second level gradient boosting was the overall best classifier. Although this type of cross validation based analysis is often used with the stacking, one can argue that evaluation data is also used in training process. This could generate a slight risk of favoring over-learning. Our other analysis, that uses a separate evaluation set, corrects this flaw.

We also evaluated our methods with CAFA3 evaluation set. This evaluation has the following benefits: *A*) Evaluation dataset was totally excluded from the cross validation training steps. *B*) Evaluation dataset allows comparison against CAFA competition results and *C*) We can use both Term Centric and Protein Centric evaluation. Here our results especially show that:

- Taxonomy and IPS location based features again improve results
- XGB and FM give again the best performance on first level
- Second level classifiers improve clearly results
- Our best methods outperform all CAFA competition methods in BP and MF results, with three or four evaluation metrics.

Our overall results with CAFA evaluation datasets show how results improve when new feature sets, less used classifiers and classifier stacking are used in the process. Improvement was strongest in CC and BP. Notice how the ranking in CC, generated by TC-AUC and *S_min_*, moves from lower ranks to top-5 for selected second level classifiers. In addition, the ranks in the first level BP results are between 2*^nd^* and 14*^th^* (from TC-AUC, *S_min_, nS_min_* and wF_max_). However, all these metrics rank our stacking methods as first or second. Similar but slightly weaker improvement is also seen in MF results.

In addition, we performed an initial third level stacking experiment similar to the second level experiment, but we abandoned it since the results did no show noticeable performance increase. Furthermore, the total computation time was considerably increased, which limits the practical application (data not shown).

### 4.2 Evaluation metrics and evaluation datasets

Definition of training and evaluation datasets is challenging for AFP problem [15, 18]. We used two quite different datasets in our analysis. Our in-house dataset has very reliable and detailed annotations for proteins. This lessens the problems caused by false negatives and false positives that often occur in AFP evaluation.

The CAFA3 dataset, on the other hand, is a well known standard dataset in AFP literature that contains a separate evaluation dataset. This allows the comparison against the methods that participated CAFA3 competition and also against the articles that have used the same dataset. Here it is critical to remember that the used training dataset must use database versions prior to CAFA3 competition.

Most AFP articles use only *F_max_* and/or *S_min_* results while comparing against the CAFA3 results. We took a different approach and used all five evaluation metrics, distributed by CAFA3 competition. This gives us more robust view. Here it is interesting to see that all other metrics, except *F_max_*, rank our second level methods very high in the results. This disagreement can be explained by the unsuitability of Protein-Centric *F_max_* for the evaluation of AFP methods [8, 18, 3]. Notice that if we would have limited analysis only to, say *S_min_* and TC-AUC, our results from CC data would look significantly better than what they look now. Or by looking just *F_max_* and TC-AUC our results from MF our results are not that impressive. These two observations point the ways how one could tweak method performance in scientific publications and underline the importance of using many metrics in parallel.

Our results for CC subset are quite unstable. Results with *S_min_* and TC-AUC are reasonably good for second-level XGB and r.mean, but other three metrics give quite weak results. Our potential explanation for this is that the CAFA CC evaluation dataset is weaker in quality than MF and BP evaluation datasets. This is supported by lower reported annotation depth in CC than in BP and MF [26]. Lower depth can lead to false negative cases in evaluation data, as the data lacks the correct classification to deeper classes in GO hierarchy. This in turn causes a perfectly correct (biologically reasonable) deep prediction to get weaker score. Still, this also signals that our system is by no means perfect, as some CAFA3 competition methods can clearly generate good performance on those metrics.

### 4.3 Future improvements

Our process can be clearly improved. Cross validation is critical component in stacking process. Unfortunately, our current process required a separate stratified cross validation for each GO class. This makes it impossible to use methods that predict multiple classes simultaneously in our stacking process. Here, we are currently developing solutions that allow stratified multi-class cross validation with large GO datasets [21].

Notice that our performance was obtained without any additional data, like overall sequence similarities, gene expression data, proteinprotein interaction data. Therefore, we argue that it should be easy to further improve this performance, either by adding these other data sources or by taking the hierarchical structure of classes into account.

## Acknowledgement

This work was funded by NNF20OC0065157 of the Novo Nordisk Foundation. Computations were partly done using resources in Biocenter Finland’s Bioinformatics platform. The authors wish to thank the Finnish Computing Competence Infrastructure (FCCI) for supporting this project with computational and data storage resources.

## Supplementary text

### 1 Introduction

This text collects text from our manuscript, *Optimizing InterProScan feature processing generates a surprisingly good protein function prediction method*, that did not fit to our main text.

### 2 Previous Research

AFP methods are a very actively studied topic, as can be seen from the reviews [5, 10] and AFP method competitions [19]. However, there has not been an extensive comparison of IPS feature preprocessing methods in previous research. Usually, the features are simply binarized, as can be seen from Table 1. We reviewed several other articles that have used IPS features. Unfortunately, it was often quite unclear how these features were actually processed. Here, our aim is to see if additional information, like e-value for the sequence feature, is useful for the prediction process.

Combining AFP predictions from different classifiers is often used to increase prediction performance. This is usually done by just pooling all the predictions [8, 12], or using a weighting scheme in pooling for increased performance [18]. Usually, the pooling is optimized over all classes simultaneously, i.e., the same classifier weight distribution is used in pooling for all classes. Here, we propose optimizing the combination of the first level predictions separately for each GO class.

Finally, we discovered an article similar to our work, while we were finalizing our manuscript [17], where different classifiers are compared in class specific prediction settings. However, that article uses smaller data, limiting analysis only to some bacterial genomes. It also lacks non-linear second level classifiers and does not compare the methods to CAFA competition results. Furthermore, this article does not discuss processing of IPS features.

**Table 1:** This table demonstrates how some AFP researchers have used IPS features. Notice that GOA uses curated links between IPS features and GO terms.

| Method | GO prediction method | Feature preprocessing | Reference |
| --- | --- | --- | --- |
| GOA database | Curated links from interpro2go | Binarized | [9] |
| GOlabeler | Logistic regression | IPS features are binarized | [18] |
| INGA2 | Cluster linked to enriched GO terms | Presumably binarized | [12] |
| Turku NLP team | Neural networks and random forest | Features first linked to GO classes. These used as input features | [8] |

### 3 Overview on used classifier methods

Table 2 represents links, for each classifier, to the original method publications and implementations.

**Table 2:** Overview of the classification algorithms used in the experiments. LR and XGB-ltr are used only in the second level classification experiments.

| Classifier name | Acronym | Reference | Implementation | Level |
| --- | --- | --- | --- | --- |
| Gradient boosting | XGB | [6] | [1] | 1 & 2 |
| Gradient boosting (LTR loss) | XGB-ltr | [6] | [1] | 2 |
| Artificial neural network | ANN | - | [11] | 1 & 2 |
| Logistic regression (elasticnet) | ELASTICNET | [20] | [11] | 1 |
| Logistic regression (l2) | LR | - | [11] | 2 |
| Support vector machine | SVM | [3] | [11] | 1 |
| Factorization machine | FM | [14] | [15] | 1 |

### 4 Details of datasets

Here, we present more detailed information about the datasets that we use in the experiments. Tables 3 and 4 show the sizes of the two used GO datasets and the table 5 shows the sizes of featuresets for the in house data.

**Table 3:** Properties of the in-house GO subset target set. Min and Max represent the minimum and maximum class sizes of the datasets. Columns 25%, 50% and 75% are the corresponding percentiles of the class sizes. Last column shows the number of classifier models, generated in five-fold cross validation

| Data | Proteins | GO classes | Density | GO class sizes |  |  |  |  | models |
| --- | --- | --- | --- | --- | --- | --- | --- | --- | --- |
|  |  |  |  | Min | 25% | 50% | 75% | Max |  |
| CC | 577424 | 1688 | 0.0077 | 5 | 66 | 225 | 891 | 577410 | 8440 |
| MF | 637552 | 3452 | 0.0028 | 11 | 61 | 150 | 498 | 637533 | 17260 |
| BP | 666349 | 11288 | 0.0028 | 4 | 41 | 123 | 493 | 666338 | 56440 |

**Table 4:** Details of sequence features in the CAFA3 dataset

| Training set |  |  |  |  |  | Evaluation set |
| --- | --- | --- | --- | --- | --- | --- |
| Data | Proteins | Labels | *e-value*<br>features | *taxonomy*<br>features | *location*<br>features | proteins |
| CC | 234604 | 1606 | 25538 | 26172 | 105524 | 1097 |
| MF | 346251 | 3241 | 29301 | 30050 | 120855 | 1101 |
| BP | 370779 | 11336 | 31417 | 32220 | 129374 | 2145 |

**Table 5:** The number of input features of feature sets in the in-house dataset.

| Data | CC | MF | BP |
| --- | --- | --- | --- |
| *e-value* | 32149 | 33737 | 35934 |
| *binary* | 32169 | 33757 | 35955 |
| *cluster* | 33289 | 34916 | 37145 |
| *taxonomy* | 32929 | 34598 | 36848 |
| *count* | 65478 | 68693 | 73121 |
| *location* | 132096 | 138565 | 147454 |

### 5 Cross validation with in-house data

The experiments with the in-house data were evaluated using cross validation. Cross validation is a standard model evaluation procedure based on testing models on data points excluded from training data. The used dataset is randomly split into *k* non-overlapping subsets (folds), and the model is trained *k* times so that one fold at a time is left out of the training process and used as an evaluation data. The *k* sets of predictions, each corresponding to one fold, are combined and evaluated with an evaluation metric to get the final performance of the model.

Training and evaluating AFP methods is challenging, as many GO classes have only few confirmed members. Here we include classes with just ten members to our prediction task. Although the very small classes are really hard to predict, they are also the most informative classes, presenting very detailed biological information. These smallest classes have a very extreme *class imbalance*, where we have over > 20 000 negative data points for one positive data point. This class imbalance is difficult with normal cross validation that uses random data splits. We looked at the existing multilabel stratified cross validation algorithms, but they were not suitable for extremely imbalanced data.

Therefore, we decided to use stratified five-fold cross validation separately for each class. Here the positive and the negative data points, of the selected class, are evenly distributed between data subsets. This ensures that each training and evaluation dataset contains both positive and negative data points, allowing precise class specific evaluation of the results. It is also important for our classifier stacking process, as it uses results from stacking as intermediate input. The drawback is that as the cross validation folds are different for each class, the classes have to be analyzed separately. This excludes multilabel classifiers, like popular Deep Neural-Networks or our current PANNZER method [16], from our comparison.

Class-specific cross validation also excludes protein-centric evaluations (explained in section Evaluation Metrics 5.1) with this data. However, it should be noted that our tests with CAFA3 evaluation datasets represent a very detailed protein-centric evaluation with four evaluation metrics. Finally, we point that separate stratified cross validation for each GO class is problematic, and our Discussion (see main text) points our future directions to solve this.

#### 5.1 Evaluation metrics

Selecting evaluation metrics for AFP is a difficult and often overlooked task [13, 10]. Still, it has a drastic impact on results, and some popular evaluation metrics are not well suited for AFP task [13, 2, 7]. Here we used metrics that are either well suited to AFP evaluation or allow comparison against the latest CAFA competition.

The evaluation metrics, used here, can be divided to *Term Centric* (TC) and *Protein Centric* (PrC) evaluation metrics. TC evaluation metrics process predictions for each GO class (or GO term) separately, generate a score for each class and then combine the separate scores by averaging them. Usually, only classes with ten or more class members are taken into analysis. PrC evaluation metrics roughly process the predictions for each protein sequence separately, generate a score for each protein and combine the generated scores by averaging them. More details on these metrics and other alternatives can be found in earlier articles [13, 19].

The evaluation metric used in our cross validation comparisons is the Term Centric Area Under the Precision Recall Curve (TC-AUCPR). It summarizes the precision-recall curve, of a GO class, as a mean of precisions weighted by the increase in recall at each threshold. Next, these class-specific AUCPR values are averaged. TC-AUCPR was selected here for its suitability for highly imbalanced data [4]. It also showed good performance in the evaluation metric comparisons [13].

We used many evaluation metrics in parallel in CAFA3 evaluation. We included the two main CAFA3 evaluation metrics: minimum semantic distance (*S_min_*) and the maximum of F-measure (*F_max_*). We also included three less used evaluation metrics: *nS_min_* (normalized *S_min_*), wF_max_ (weighted *F_max_*) and the Term Centric area under Receiver Operating Characteristics curve (TC-AUC). All these measures, except for TC-AUC, are PrC metrics. CAFA3 competition organizers distribute the results for all CAFA3 competition participants with all these five metrics.

Using these five evaluation metrics in parallel has its benefits: Each metric is expected to have its own biases and errors. The effect of these weaknesses is lessened when we monitor five metrics in parallel. Furthermore, we and others have shown that *F_max_*, the main evaluation metric in CAFA3 competition, is a biased metric [13, 7, 2]. It favors methods that predict GO classes very close to the root, or the root of the GO structure. Here the inclusion of other evaluation metrics, *S_min_, nS_min_*, wF_max_ and TC-AUC allows us to check if they can generate a more reliable consensus.

## Notes

http://ekhidna2.biocenter.helsinki.fi/AFP

### Competing Interest Statement

The authors have declared no competing interest.

### Summary of Updates

Supplementary text added at the end of the manuscript

